# Characterising a stable five-species microbial community for use in experimental evolution and ecology

**DOI:** 10.1101/2020.04.24.059097

**Authors:** Meaghan Castledine, Joseph Pennycook, Arthur Newbury, Luke Lear, Zoltan Erdos, Rai Lewis, Suzanne Kay, Dirk Sanders, David Sünderhauf, Angus Buckling, Elze Hesse, Daniel Padfield

## Abstract

Model microbial communities are regularly used to test ecological and evolutionary theory as they are easy to manipulate and have fast generation times, allowing for large-scale, high throughput experiments. A key assumption for most model microbial communities is that they stably coexist, but this is rarely tested experimentally. Here we report the (dis)assembly of a five-species microbial community from a metacommunity of soil microbes that can be used for future experiments. Using reciprocal invasion from rare experiments we show that all species can coexist and we demonstrate that the community is stable for a long time (∼600 generations). Crucially for future work, we show that each species can be identified by their plate morphologies, even after >1 year in co-culture. We characterise pairwise species interactions and produce high-quality reference genomes for each species. This stable five-species community can be used to test key questions in microbial ecology and evolution.

## Introduction

Controlled experiments on synthetic microbial communities have been used to test general ecological and evolutionary theories about - for instance - species coexistence [1], adaptive radiations [2–4], and ecosystem functioning [5, 6]. The short generation times and large population sizes of bacteria make them ideal for testing ideas that are experimentally intractable in other model systems. Culturable bacteria are also easy to manipulate, allowing us to change species richness [5, 7, 8], abundances [9, 10], and even interaction type [11, 12] and strength [13] in a systematic and increasingly high throughput way.

The majority of previous experimental work has been done on monocultures or in synthetic communities consisting of only a few species. However, an increasing number of studies have used more diverse microbial communities to study how species rich communities assemble and their stability, to understand the maintenance of the high levels of microbial diversity seen in nature [8, 14–17]. Experiments on both simple and complex model communities are key to understanding the processes underpinning community assembly and coexistence, and there is an inherent trade-off between more detailed mechanistic understanding and complexity [18].

A key assumption of both approaches is that the community members stably coexist, meaning that the densities of species in the system do not show long-term trends [19]. Modern coexistence theory presents the ability to invade from rare (negative frequency dependence) which tests that each species is able to increase from low density when in the presence of the rest of the community, as a key test of coexistence [19, 20]. Experimentally establishing that model communities are stable and that species coexist is key to increasing the likelihood that the findings are relevant to natural communities (where high levels of diversity are maintained, composition is relatively stable through time across space [21–23], and where patterns of diversity cannot be explained solely through neutral processes [24–26]). Second, when a model community is not stable, it risks biasing our understanding of species interactions. Specifically, species will be driven locally extinct in unstable communities, which will necessarily bias interactions towards being negative. Third, demonstrating stable coexistence and understanding the mechanisms of coexistence (e.g. equalising or stabilising [19]) would increase the generalisability of findings and allow future work to understand under what circumstances stability breaks down.

There are several proposed methods to measure invasion growth rate [27], but these become more complicated to test as species diversity increases, where you have to test whether each species has a positive invasion growth rate while it is absent and the resident community is at equilibrium [28]. For example, following the removal of a species, the resident community may destabilise or convert to an alternative stable state [29]. Explicit tests of coexistence in model microbial communities have been done when communities are simple and contain only strains of one species [30–32], two [33], or three species [7], but they remain scarce when species diversity goes beyond this ([5, 8, 13, 15, 17], but see [34]). Most model microbial communities are performed in batch culture, where populations are regularly transferred into fresh microcosms [2], meaning that “equilibrium” is never reached in a measurable sense. Consequently, microbiologists have used invasion-from-rare assays to test for negative frequency dependence and stable coexistence. In these assays, the community is inoculated into fresh media, but one species (the invader) is at 100-fold lower density than the other species (the resident community). If each species has a higher growth rate when rare, relative to the resident community (a relative invader fitness greater than 1) then the community shows negative frequency dependence and is likely to be stable [31, 35, 36].

Here, we report the characterisation of a stable 5 species community that can be used for future experiments in ecology and evolution. We cultured a community pool of 46 isolates retrieved from soil over several weeks in lab media and the final community composition was determined. As outcomes were highly repeatable, we took a single replicate community and conducted reciprocal invasion experiments to check coexistence. In addition, we measured species interactions using spent media and co-culture, tracked long-term community dynamics, and generated high quality reference genomes. Finally, we discuss potential questions that could be answered with this - or any - stable community when so much information about its constituent species is known.

## Methods

### Community disassembly experiment

We created a model microbial community consisting of 5 species that (dis)assembled for approximately 85 generations (Figure 1a). Briefly, this community was formed by inoculating 46 bacterial strains – belonging to 27 genera (Table S1) – that were isolated from an experimental compost community [37] into 6mL growth medium (1/64 Tryptic Soy Broth (TSB) diluted with demineralised HLJO) in 25mL glass vials with loosened plastic lids. Forty eight cultures were set up, incubated statically for 13 weeks at 28°C. For the first five weeks no transfers were made, after which weekly transfers were done, passaging 1% (60µl) of each community into fresh 6mL microcosms every week. After this time, the 48 communities converged on a similar composition of ∼5 dominant species. We selected a single replicate of these to explore its stability.

**Figure 1.**
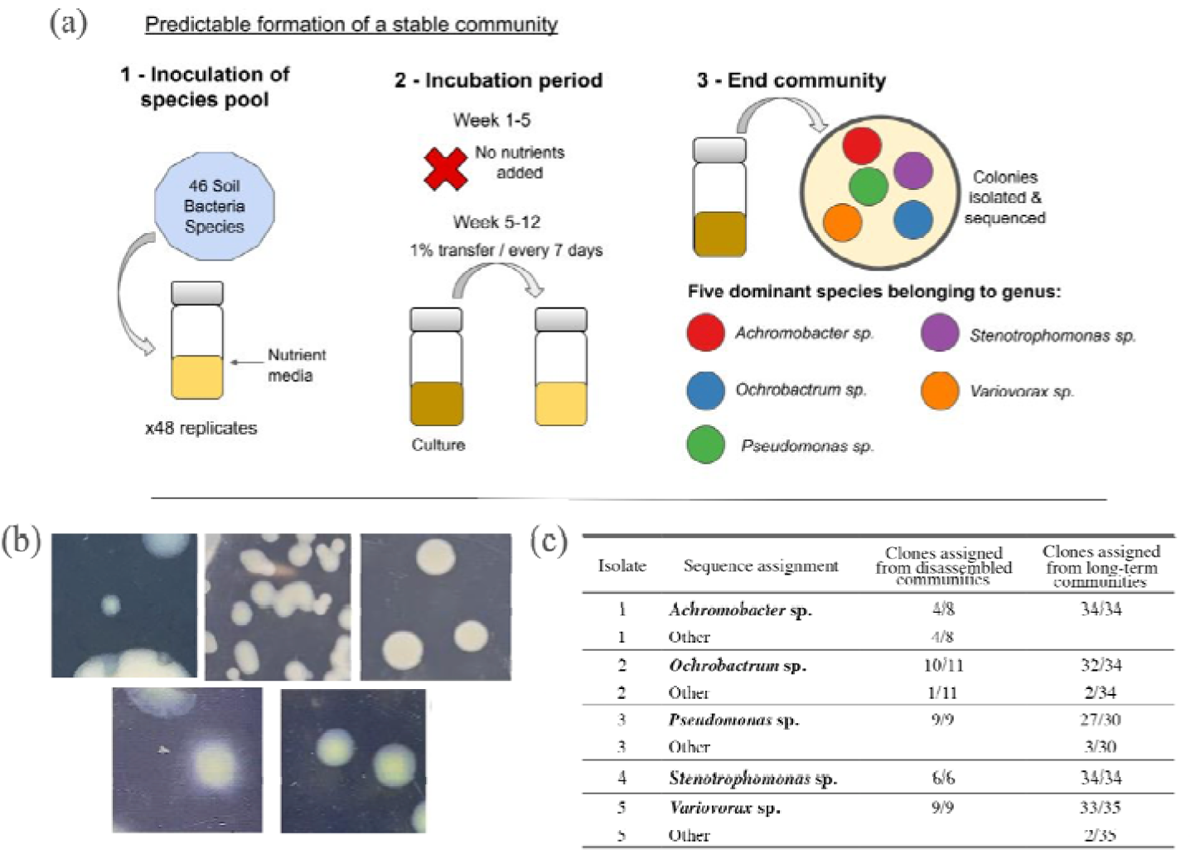
Community (dis)assembly experiment to form a stable community. (a) Schematic of the experimental design for the formation of the stable microbial community. (b) Five morphs dominated the community. (1) *Achromobacter* sp., the smallest colony, circular and uniform in size. (2) *Ochrobactrum* sp. white/grey in colour and opaque. (3) *Pseudomonas* sp. is the largest colony, cream in colour and opaque. (4) *Stenotrophomonas* sp. is typically larger than *Achromobacter* sp. although variable in size and has a stronger colour. (5) *Variovorax* sp. has a spreading mat surrounding the colony and is variable in size. (c) Reliability of morphotypes for identifying the different species.

This community consisted of five species, *Achromobacter* sp., *Ochrobactrum* sp., *Pseudomonas* sp., *Stenotrophomonas* sp., and *Variovorax* sp.. Crucially, these species could be reliably identified by their colony morphology (Figure 1b), allowing us to track each species in the community and re-isolate them for phenotyping and clone re-sequencing. A single clone of each member of this community was picked and grown in monoculture for two days in 6mL of 1/64 TSB, shaking, in 25mL glass vials with loosened plastic lids at 28°C to achieve high cell densities. These samples were then cryogenically frozen at −70°C in glycerol (final concentration: 25%). These clones were used to recreate the community for subsequent experiments.

### Sanger sequencing of 16S rRNA of each morphotype

To understand how well our characterisation of different morphotypes represent different species, we sequenced the 16S rRNA gene of multiple replicates of each morphotype. Of the 5 distinct morphotypes, we picked at least 8 colonies across all 48 microcosms. Polymerase chain reactions (PCRs) were performed in 20µL reactions containing 10µL of DreamTaq Green PCR MasterMix (2X) (Thermo Scientific), 0.4µL each of the 338F and 518R primers (10µM) and 2µL of 1:100 diluted culture. In total 51 samples were Sanger sequenced using the 515F primer (Eurofins Genomics).

The Sanger sequencing files were analysed in *R*, with *R* functions specified as *Rpackage::function*. Chromatograms for each sample were created using *sangeranalyseR::secondary.peaks* [38]. Low-quality bases were trimmed from the start and end of each file using *sangeranalyseR::trimm.mott* which uses Mott’s modified trimming algorithm, with a cut-off of 40 to define a bad quality score. Files were further filtered to retain only sequences that were longer than 100 base pairs long, had five or less secondary peaks (where there is more than one peak at a given position; a secondary peak was defined when the lower peak was at least a third as high as the higher peak), and had a mean quality score of >30. This left 43 trimmed and cleaned sequences that went forward to further analysis.

For each of the 5 identified morphs, we calculated a pairwise distance matrix from the DNA sequences using *ape::dist.dna* [39], with a phylogenetic tree being constructed and plotted using *ape::njs* and *ape::plot.phylo*. If all replicates of the same morph were attributed to the same level on the tree, a single consensus sequence for the morph was constructed using *msa::msaConsensusSequence* [40] after aligning multiple sequences of the same morph using *msa::msa*. In instances where there was more than a single branch of the tree, samples within a morph were split along branches to create multiple consensus sequences. Consensus sequences were aligned using *msa::msa* to look at similarities between morphs. Taxonomy was assigned to genus level using *dada2::assignTaxonomy* [41], which implements the RDP Naive Bayesian Classifier to assign taxonomy to the RDP database (v16) [42].

### Invasion-from-rare assays

To explicitly test for coexistence, we conducted invasion-from-rare assays [4]. To assess whether all species were necessary for coexistence, we did assays where each species was inoculated from rare in every species by diversity combination. In total, this yielded six replicates of 75 treatments. Species were grown from frozen to achieve high cell densities as described above. Cell density (colony-forming units [CFU]) was estimated from optical density (600 nm wavelength; OD_600_) using a species-specific calibration curve (Table S3 for equations) and normalised to 10^5^ CFU μL^-1^. Focal species were invaded at a 100-fold lower density (1.6µL) than that of the resident community (160µL in total). When there was more than one resident species, resident communities were created from equal ratios of each species. Cultures were incubated for one week, after which samples were cryogenically frozen at −70°C in glycerol. Species abundances (CFU mL^-1^) were estimated by plating cultures onto KB (King’s medium B) agar and incubating for two days at 28°C.

Relative invader growth rate of the focal species was calculated as the ratio of estimated Malthusian parameters (*m*), *m*_focal_: *m*_community_ where *m*_community_ is the total density change of all the other populations combined. *m* = *ln(N_1_ / N_0_)/t* where *N_1_* is the final density, *N_0_* is starting density, and *t* is the assay time (7 days) [4]. We performed independent one-sample t-tests on each combination of focal species by resident community combination, with the null-hypothesis being that the mean relative invader growth rate equals 1. A relative invader growth rate above 1 would indicate that the invader had a faster growth rate than the resident community, demonstrating negative frequency dependence. This resulted in 75 different statistical tests; *P* values were adjusted using the false discovery rate (*fdr*) method [43].

### Medium-term persistence assays

To test whether diversity impacts coexistence and persistence, and to quantify pairwise and indirect (higher order) interactions, each species was also inoculated at equal density in every possible community combination, from monoculture to five-species. Species monocultures were prepared with 20μL of each species inoculated (total inoculum varied across diversity levels) into fresh microcosms for each community combination. Twelve replicates of each unique community combination were then incubated for either 1, 2, 3 or 4 weeks (three replicates each) without resource replenishment. These assays allowed time for extinctions to occur, to measure the effect of resource scarcity and secondary metabolites for cultures left for 3 or 4 weeks, and for us to calculate species interactions of the community. At the end of each incubation period, individual species abundance (CFU mL^-1^) was estimated using methods described above. Abundances from week 1 were removed as it was visually clear that equilibrium density had not been reached for most species.

We calculated pairwise species interactions (e.g. effect of species *2* on species *1*, *w_1|2_*) as the abundance of species *1* in the presence of species *2* relative to that of species *1* grown in monoculture (no interspecific interactions): *w_1|2_ = N_1|2_/N_1_*. If *w* is greater than 1, then the interaction is positive, whereas a *w* < 1 demonstrates competitive interactions, with a value of 0 demonstrating competitive exclusion. To get a broad perspective of the nature of species interactions, we calculated the average (the mean of ∼9 replicates) for each pairwise species interaction.

We estimated indirect (higher order) interactions from the persistence assays using a modified version of a framework originally developed to help quantify higher order antibiotic interactions on *Escherichia coli* [44]. As our assays cannot determine whether the presence of a third, fourth, or fifth species changes competitor density (an interaction chain) or changes per capita competitive effects (higher order interaction), our method calculates indirect interactions generally. This method overcomes the issues of the additive model when single species have large impacts on fitness, such that the additive expectation can never be achieved (i.e. a population size < 0)[45, 46].

For each community combination with more than two species (e.g. species *1, 2, 3*), the predicted effect on the focal species, ŵ*_1|23_*, is the product of the independent pairwise interactions: ŵ*_1|23_ = w_1|2_ x w_1|3_*. The indirect interaction (*ii*) of any community combination on focal species *1*, *ii*_1|23_, was calculated as the deviation of the observed abundance relative to monoculture, *w*_1|23_, from that predicted from pairwise interactions: *ii_1|23_ =* ŵ*_1|23_ - w_1|23_*. Using this definition, if an *ii* is the same sign as the original prediction (ŵ*_1|23_*), then it indicates synergism: the combined presence of species 2 and 3 on species 1 is greater than either on their own. If an *ii* is the opposite sign of the original prediction, then it indicates non-additivity and potential buffering: the combined presence of species 2 and 3 on species 1 is less than either on their own. To make all synergistic indirect interactions positive (and buffering interactions negative), we multiplied indirect interactions of predicted negative interactions by −1. We looked at whether the type of indirect interaction (buffering or synergism) was related to the strength of the expected pairwise interaction using a linear model with expected pairwise interaction as the response and type of indirect interaction as the response. This was only done on predicted negative interactions as there were only seven predicted positive interactions (all belonging to *Variovorax* sp.) in the dataset.

The original Tekin method rescales interaction estimates and quantifies indirect interactions after accounting for all interactions at lower species combinations (for example, in four species communities the indirect interaction of *ii_1|234_* includes taking into account the estimate of *ii_1|23_*, *ii_1|24_*, and *ii_1|34_*). Recent work has discussed whether pairwise interactions alone can predict species abundances, so we chose to calculate all indirect interactions relative to the prediction of only the pairwise interaction estimates.

### Supernatant assays

Cell-free supernatants were used to establish cross-feeding assays in 96-well plates. To obtain the supernatants, three replicate populations per species were established by inoculating a single clone in 6mL of 1/64 TSB in 30mL glass vials. Following 2 days of growth at 28°C, we centrifuged individual cultures at 3000 rpm for 10 minutes, and filtered these through 0.22mm syringe filters (Merck, Millex PES). We confirmed supernatants were cell free by plating these onto KB agar. For each focal strain we inoculated ∼5 x 10^4^ cells from an overnight culture in triplicate wells, each containing either 200µL of fresh 1/64 TSB or cell-free supernatant of either the same or a different species (n=3 per pairwise combination). Following 48 hours of growth at 28°C, serial-diluted cultures were plated onto KB agar to obtain final cell densities.

Growth rate per day was calculated as *ln(N_1_ / N_0_)/t*, where *N_1_* is final density, *N_0_* is the initial density, and *t* is two (the number of days of growth). If the growth rate was negative, we replaced the value with 0.001 to prevent negative values of relative growth required for comparisons between species. Relative growth rate was calculated by dividing the growth rate in supernatant by the growth rate in fresh media. In this way, if the value is above 1, the supernatant has a positive effect on growth, and if it is below 1 it has a negative effect on growth. This method gives a measure of the intraspecific effect and interspecific effect of each species by comparing the relative growth in each species’ own supernatant to those in other species’. No formal statistics were done on the interactions of individual species, choosing instead to only describe the types and strengths of interactions. These measurements may help explain why we see coexistence, and can act as a benchmark against which to understand how interactions change after future experiments.

We looked at whether estimates of species interactions from supernatant assays and co-culture experiments matched both qualitatively and quantitatively. Quantitatively, we used standardised major axis (SMA) regression to look at the relationship between the two estimates for each pairwise combination. SMA was useful here as we have no expectation about one variable being the response or the predictor, and allows us to test for an expected slope of 1. Qualitatively, we looked at whether the same sign of interaction (positive or negative) occurred when calculating pairwise interactions using co-culture and supernatant assays.

### Long-term stability

To understand long-term stability of the five-species community, 12 cultures were transferred weekly for 60 weeks. To start the cultures, monocultures of species were grown from the freezer stocks to achieve high cell densities as described above. Populations were set-up using 20µL inoculum of each species’ culture normalised using the species-specific calibration curves and incubated at 28°C [47]. Serial 100-fold dilutions (60μL culture into 6mL 1/64 TSB as described earlier) took place every week for over a year. Cultures were plated every two weeks to check for contamination. After 60 weeks (at least 400 generations), our plate counts and identification of morphotypes indicated that all 5 species remained present in all cultures. To confirm this, we sequenced the 16S rRNA gene of multiple replicates of each morphotype (species) across multiple replicates. We picked at least 30 colonies of each morphotype across all 12 populations, with 169 samples in total. DNA was extracted by following Qiagen’s standard DNeasy UltraClean Microbial protocol, and the concentration of gDNA in each sample was determined using QuBit. A conserved fragment from the hypervariable region of the 16s rRNA gene was targeted using N501F and N806R primers and amplicon sequencing was undertaken by the Centre of Genomic Research (Liverpool, UK) using an Illumina MiSeq to create 2×250bp paired end reads.

We processed and analysed the sequence data in R using the packages *dada2* and *phyloseq* [48]. Following the standard full stack workflow, we estimated error rates, inferred and merged sequences, constructed a sequence table, removed chimeric sequences and assigned taxonomy. During processing, forward and reverse reads were truncated between 25 and 250 nucleotide positions due to poor quality scores. Assembled amplicon sequence variants (ASVs) were assigned taxonomy using the RDP database. We then aggregated OTUs at the genus level as we knew each of the five species belonged to a different genus. One sample was removed as it only had 135 reads, and one sample was removed because the dominant ASV (the identity of the clone) was <0.95, indicating that the colony likely consisted of multiple species.

### Whole genome sequencing of each morphotype

To better understand the genomic features of each species, we did *de novo* hybrid assembly using both short- and long-read sequencing. DNA was using a NEB monarch genomic DNA kit following the manufacturer’s instructions, apart from for *Ochrobactrum* spp. samples, which were vortexed with beads and had an extra proteinase-K treatment to aid lysis. Short read Illumina sequencing was done by the Exeter Sequencing Service on a HiSeq 2500 ran in standard mode. Long read sequencing was done using the PacBio HiFi platform on a Sequell II using one SMRT cell by the Centre of Genomic Research (Liverpool, UK) and quality processed by them.

Hybrid assembly was done using *HybridSPAdes* [49] with the option “*-isolate*” specified. The assemblies were polished with the trimmed Illumina data using *Pilon* [50] under default settings. *Circlator* [51] was used to circularise the assemblies, which utilised the PacBio reads which were corrected using *Canu* [52] with the option “*genomeSize*” set to the approximate size for each isolate (estimated from the BLAST results of each isolates 16S sanger sequence). All contigs <500bp were removed, and a contig that was identified as the phage *phiX* - used as spike in for the short-read sequencing - was removed. Assemblies were annotated using *prokka* [53] with default settings. We ran *breseq* [54] to map the short reads back to each assembly to identify potential errors in the assembly and to estimate depth.

Taxonomy was estimated using *gtdbtk* [55] which uses FastANI [56] to do whole genome alignment, and by using *dada2* to assign taxonomy using 16S genes in the assemblies. The quality of the assemblies was assessed using *CheckM2* [57].

## Results

### Final (dis)assembled community

From plating all 48 communities, we identified 5 distinct morphotypes that were dominant in almost all communities. Based on Sanger sequencing, the five colony morphotypes were identified as *Achromobacter* sp., *Ochrobactrum* sp., *Pseudomonas* sp., *Stenotrophomonas* sp., and *Variovorax* sp. (Figure 1b). Overall we have high confidence that these morphs represent different species. For *Pseudomonas* sp., *Stenotrophomonas* sp., and *Variovorax* sp., all morphotype replicates were assigned to the same consensus sequence. For *Ochrobactrum* sp., 11/12 morphotype replicates were aligned to a single consensus sequence, with the remaining replicate assigned to the closely related *Rhizobium* sp.. As both sequences are on the same branch of the global tree, this morph is unlikely to have been misidentified to any other of the four morphs. For isolate 1 (likely *Achromobacter* sp.), 4/8 replicates were assigned to *Achromobacter* sp., 3/8 were assigned to *Stenotrophomonas* sp. and 1 replicate was assigned to *Pseudomonas* sp. (Figure 1C). This may lead to some underreporting of *Achromobacter* sp. and overreporting of *Stenotrophomonas* sp., but it is unlikely to qualitatively alter results of experiments that use this community. We chose a single replicate community and a single isolate of each colony morphology from which to do all future experiments.

### Reciprocal invasion-from-rare experiments

We determined whether all species were key for coexistence by invading each species from rare in all species combinations. Species could invade, irrespective of resident community composition or diversity, in 71 of the 75 combinations (one-sample t-test: *P_adj_* < 0.05, Figure 2). Moreover, there was considerable variation in relative invader growth rate, ranging from a minimum of 1.12 to a maximum of 4.34. Of the non-significant combinations, three were when *Achromobacter* sp. was the invading species (including when it was invading into the other 4 species together; Figure 2a), the other when *Stenotrophomonas* sp. was invading into a community of *Achromobacter* sp., *Variovorax* sp. and *Ochrobactrum* sp. However, although not significant, mean estimates of relative invader growth rate were still above 1.

**Figure 2.**
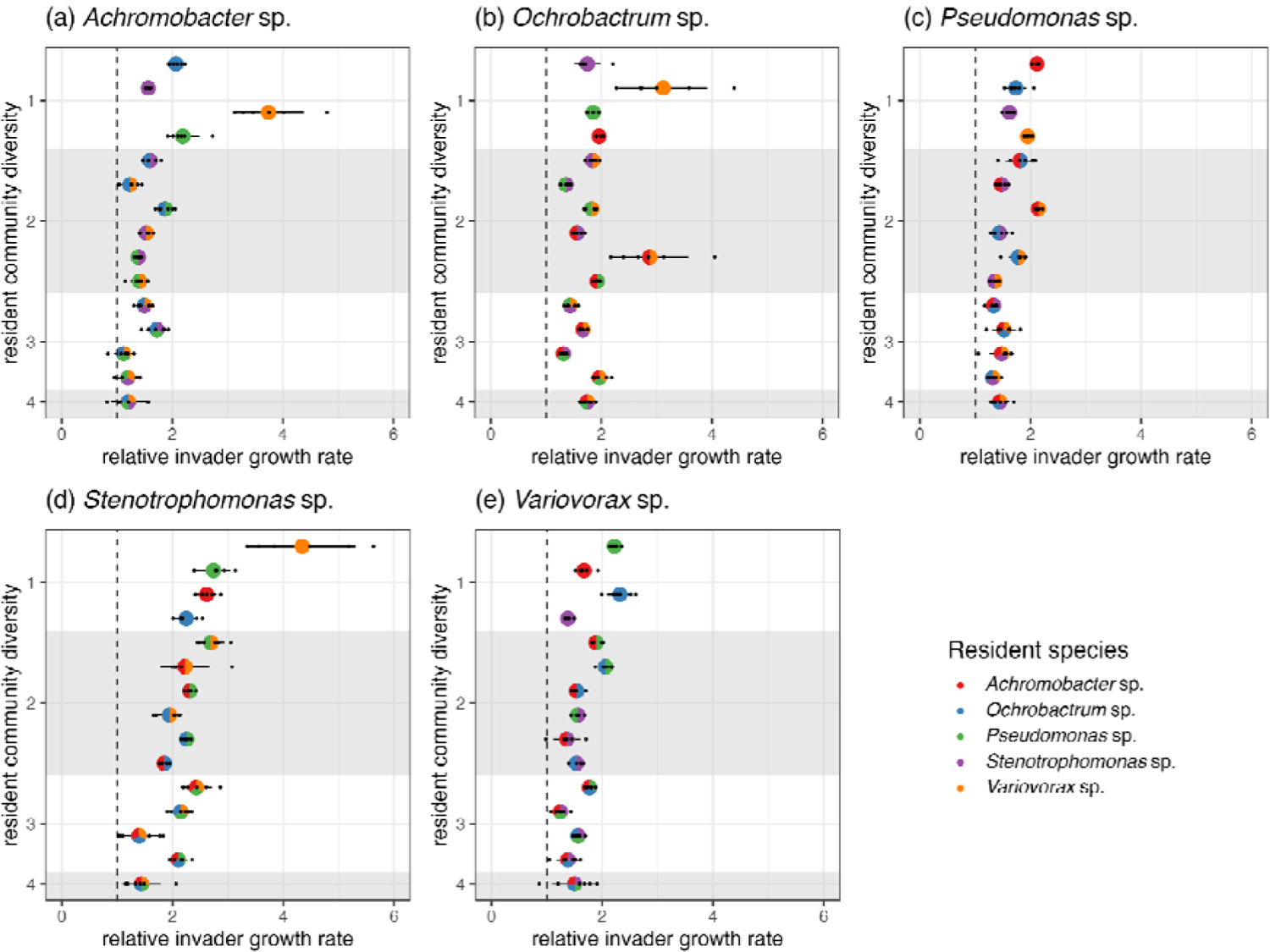
Ability of species to invade-from-rare in all combinations of species and diversity. Panels represent the invader species. Colours of pie points reflect the mix of the resident community. Black points are observed values of relative invader growth rate (n∼6), pie points are the estimate of relative invader growth rate from each one-sample t-test, and error bars represent lower and upper 95% confidence intervals. Shaded regions separate different levels of resident community diversity; dashed, vertical line drawn where relative invader growth rate = 1, values above which indicate the invading species can successfully re-establish from rare.

### Interactions based on co-culture and spent media

From doing co-culture assays in all combinations of species diversity, we were able to explore the nature of species interactions between species pairs in the model community and also how these changed at higher levels of diversity, allowing us to estimate indirect interactions. Four of the five species (*Achromobacter* sp., *Ochrobactrum* sp., *Pseudomonas* sp., *Stenotrophomonas* sp.) had competitive interactions, growing worse in co-culture with other species compared to monoculture (Figure 3b), with three of the species negatively impacted by all other species. Although *Variovorax* sp. negatively impacted the growth of all other species, it grew better in the presence of three of the other species, demonstrating asymmetric positive interactions (Figure 3b). The only species which had a symmetrically competitive interaction with *Variovorax* sp. was *Stenotrophomas* sp..

**Figure 3.**
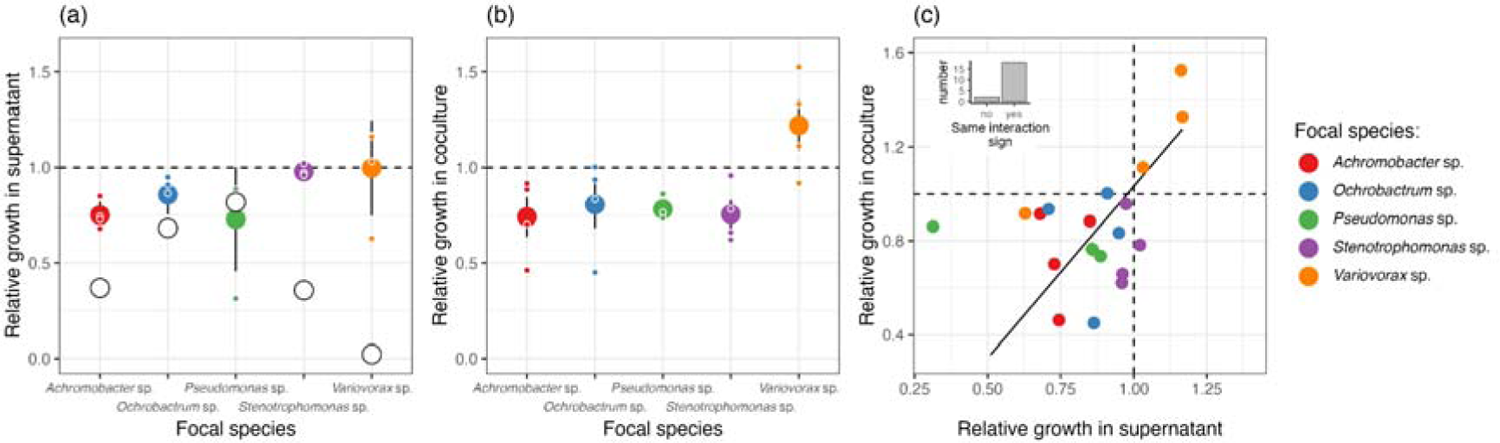
Estimated pairwise interactions of the model community from supernatant and coculture assays. (a) Growth of each species was in the supernatant of each species (including itself) and fresh media. (b) Growth of each species was measured in coculture and in monoculture. (c) Relationship between estimates of interactions estimated from supernatant assays and coculture. In all plots, values below one indicate negative interactions. In (a) and (b) large coloured points are the mean value of all interspecific interactions with 95% confidence intervals, and small coloured points represent individual interspecific interactions (the median value of three technical replicates). In (c) the solid fitted line is the best fit of a standardised major axis regression, the dashed lines demonstrate where interactions go from being negative to positive, and the inset plot demonstrates how often the interactions are qualitatively the same type using coculture and supernatant assays.

In combinations of more than 2 species, we saw large variation in estimates of indirect interactions (Figure 4a). Overall, *Ochrobactrum* sp., *Pseudomonas* sp. and *Stenotrophomas* sp. had on average negative indirect interactions, which given they had negative pairwise species interactions, indicates interactions in multispecies culture are more synergistic than would be expected by pairwise interactions alone. The opposite is true for *Achromobacter* sp., where we see on average buffering indirect interactions where interactions in multispecies culture are less than would be expected by pairwise interactions alone. *Variovorax* sp. has huge variation in indirect effects, indicating that interactions from coculture are bad at estimating how it will do in multispecies culture.

**Figure 4.**
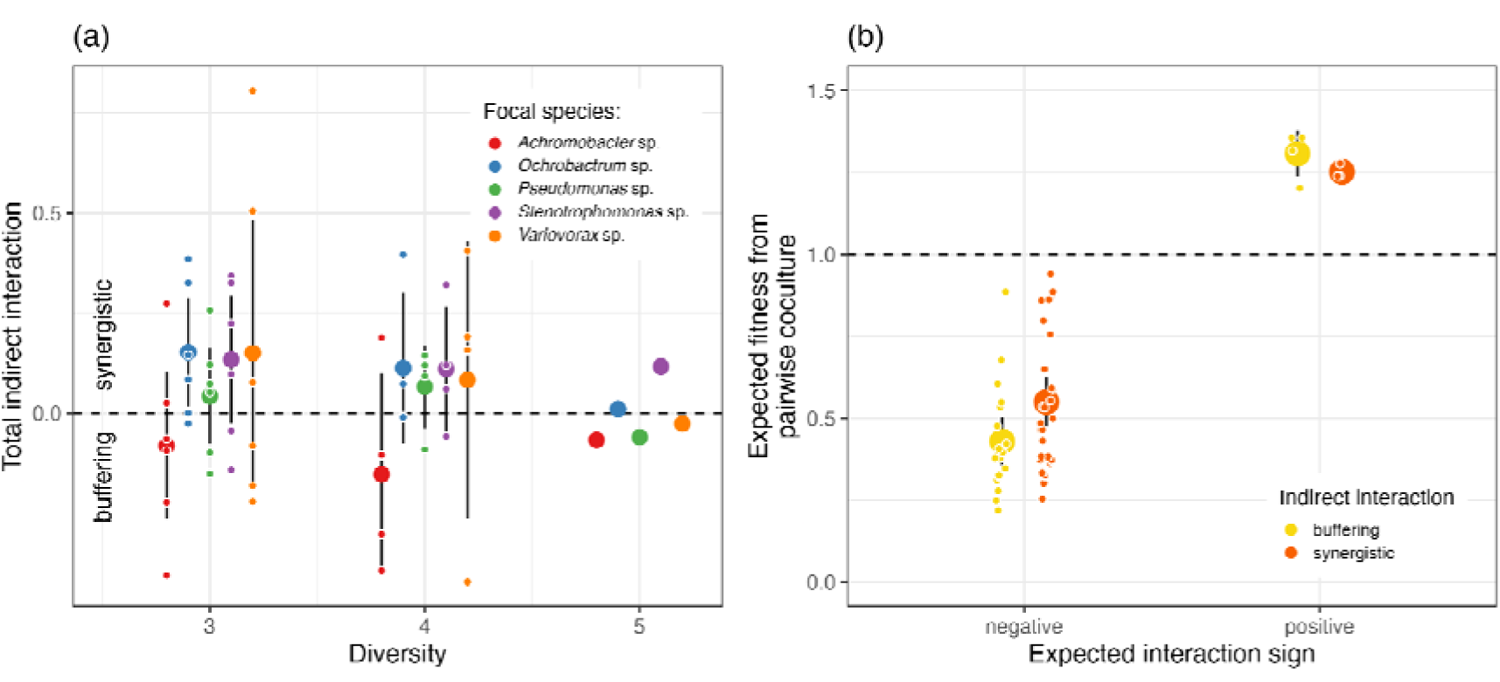
Estimated indirect interactions of the model community from coculture assays. (a) Total indirect interactions across levels of diversity show lots of variation across all focal species. (b) Impact of the strength of negative and positive pairwise interactions on whether the indirect interaction is synergistic (stronger than expected) or buffering (less than expected). As negative and positive pairwise interactions get stronger, there is a tendency for the indirect interaction to be buffering. In (a) and (b) large, coloured points are the mean value of all interspecific interactions with 95% confidence intervals, and small coloured points represent individual indirect interactions (the median value of three technical replicates).

When looking at the indirect interactions of all species together (Figure 4), we found that 41% were synergistic (23 out of 55), with 59% being buffering, when compared to the expected effect of pairwise interactions. For three- and four-species communities, buffering was found 40% of the time, and synergistic 60% of the time, with this pattern switching when all 5 species were present (albeit only with 5 data points). It is important to note that the definition of synergism and buffering of indirect interactions is sensitive to the null model we are comparing against, which we set here simply as the impact expected from pairwise interactions alone [46]. If we included the indirect interactions of three species combinations into the estimated indirect interactions of four species, we may have seen more buffering [44]. We chose not to do this for simplicity in explanation and interpretation.

No focal species consistently had buffering or synergistic indirect interactions irrespective of species combination they were co-cultured with (Figure 4a). Across all indirect interactions, we found that as pairwise interactions intensify, indirect interactions are more likely to be buffering (lower effect than expected), with synergistic indirect interactions more likely when the expected pairwise interaction is weaker (Figure 4b). This pattern is statistically significant for expected negative interactions (ANOVA between models with and without indirect interaction type as a predictor: *F_1,47_* = 4.51, *p* = 0.039), and the pattern is the same for expected positive interactions, albeit a lack of expected positive interactions prevents us from doing formal statistical analysis.

The supernatant assays allowed us to get an estimate of intraspecific competition, as well as interspecific competition, with coexistence expected when intraspecific competition is greater than interspecific competition. In four of the five species (*Pseudomonas* sp. being the exception, where no difference was detected) intraspecific competition was stronger than the interspecific effect (Figure 3a). In 3 of the species the strength of intraspecific competition was at least twice that of the estimate of the lower 95% confidence interval of interspecific competition, and in *Variovorax* sp. intraspecific competition was twelve times stronger than interspecific competition.

Both coculture and supernatant assays are commonly used to estimate species interactions, but little work has compared the two methods. We find a positive relationship between estimates of interactions from supernatant and coculture (slope = 1.46, 95%CI = 0.86-2.45, Figure 3c), but the model only explains 15% of the variation in the data, meaning that the fit between the supernatant and coculture estimates is poor. However, qualitatively, both approaches characterised the interaction as the same sign (positive or negative) in 18 of the 20 interaction estimates (Figure 3c, inset). In the two instances where they disagreed, *w_o|a_* and *w_s|o_*, one of the estimates was more or less neutral and therefore difficult to define as either positive or negative.

### Long-term coexistence

To examine whether the five-species community was stable long-term, we re-sequenced and visually identified multiple clones of each morphotype across six communities that had been cultured for 60 weeks. Crucially, this will also tell us whether morphotype differences still exist after long-term culturing which is key for the use of model communities in experimental-evolution studies.

Of the 167 samples that passed quality control, 160 (∼96%) were correctly identified from colony morphology (the genus assigned by the 16S sequencing was the same as that assigned by the assessor). Of the seven that were mis-assigned, 2 *Achromobacter* sp. were identified as *Ochrobactrum* sp., 2 *Pseudomonas* sp. were identified as *Variovorax* sp., 2 *Variovorax* sp. were identified as *Pseudomonas* sp., and 1 *Achromobacter* sp. was identified as *Pseudomonas* sp.. These mis-assignments do not match reliably with those from the initial Sanger sequencing but remain extremely rare so are again unlikely to qualitatively alter results of experiments that use this community. This confirms that the community has long-term stability and that the individual species can still be identified by their morphotype after this time.

### Genomic composition

Using short- and long-read sequencing, we created high quality reference genomes for each of the five species. All of the genomes have very high completion (>99.9%) and very low contamination (<2%), with the worst assembly having only 3 contigs (Table 1). Reassuringly, taxonomic assignment from GTDBtk corresponds to the genus-level taxon assignment from the Sanger sequencing, but for 3 of the 5 isolates a species name is now available (Table 1). Using GTDBtk to assign taxonomy using genome similarity, 4 of the species show >98% similarity to previously sequenced genomes (Table 1). The isolate genomes have now been submitted to NCBI and have been given strain names: *Achrombacter veterisilvae* AB1, *Ochrobactrum teleogrylli* AB1, *Pseudomonas fluorescens* AB1, *Stenotrophomonas sp.* AB1, and *Variovorax sp.* AB1. Genomic elements associated with plasmids, prophages, secondary metabolites, defence systems, and antibiotic resistance are present in all species. These genomes can be used as high quality references to look at genetic changes in future experimental evolution experiments.

**Table 1.** Genome characteristics and taxonomic assignment of the 5 species genomes.

| Isolate | NCBI submission name | Sanger sequence assignment | GTDBtk assignment | 16s copy number | Genome size (bp) | Number of contigs | GC content | Total coding sequences | Completeness | Contamination |
| --- | --- | --- | --- | --- | --- | --- | --- | --- | --- | --- |
| 1 | *Achromobacter* *veterisilvae* AB1 | *Achromobacter* | *Achromobacter* *veterisilvae* | 4 | 6,549,044 | 1 | 0.67 | 5,991 | 100.00 | 0.98 |
| 2 | *Ochrobactrum* *teleogrylli* AB1 | *Ochrobactrum* | *Ochrobactrum_B* *teleogrylli* | 1 | 5,064,753 | 3 | 0.57 | 4,714 | 99.97 | 1.41 |
| 3 | *Pseudomonas* *fluorescens* AB1 | *Pseudomonas* | *Pseudomonas_F* *fluorescens_BH* | 2 | 6,759,727 | 1 | 0.60 | 6,053 | 100.00 | 0.15 |
| 4 | *Stenotrophomonas* sp. AB1 (2024) | *Variovorax* | *Variovorax* sp003019815 | 1 | 6,611,336 | 2 | 0.68 | 6,041 | 99.99 | 0.37 |
| 5 | *Variovorax* sp. AB1 (2024) | *Stenotrophomonas* | *Stenotrophomonas* spp. | 1 | 3,926,254 | 2 | 0.66 | 3,384 | 100.00 | 0.04 |

## Discussion

We characterised a model five species community natural community that can be used in experiments in microbial ecology and evolution. The community contains species with mostly competitive interactions, but *Variovorax* sp. demonstrates exploitative (potentially cross-feeding) interactions with 3 of the 4 other species. We found that estimates of interactions from supernatant assays and coculture assays agreed qualitatively, but not quantitatively, with estimates of interactions in coculture being stronger on average than those in supernatant. This suggests that contact dependent interactions (e.g. T6SS and biofilm formation) may also be important, but it also may just reflect methodological differences in the approaches. We found evidence of indirect interactions, but they became weaker (and more likely to be buffering) as expected pairwise interaction increased. This meant that in the full five species community, indirect interactions were relatively weak.

This community has numerous advantages as a model system for ecological and evolutionary studies. Sequencing has confirmed that each species has a distinct colony morphology that means it can be tracked and re-isolated from the other members of the community, and that these colony morphologies remain distinct through a year of long-term culture. This makes this community ideal for experimental evolution. Furthermore, we used invasion-from-rare assays to experimentally demonstrate species coexistence, and sequencing showed that the community is stable through long-term (>1 year) batch culture. We encourage - where possible - more explicit tests of coexistence in microbial community experiments. Our model community is composed of diverse bacterial species all isolated from the same soil sample. Even so, this community is not designed to mimic a soil community.

Instead the aim was to create a community that we could show empirically to be stable that could then be used to investigate general questions about ecology and evolution. The long-term stability of the community and the unique ability to re-isolate each species to test for evolutionary responses in phenotype make the community perfect for doing long-term experimental evolution experiments, a multi-species version of Lenski’s pioneering LTEE. The genome assemblies provide a reference database to track genetic changes through evolutionary time.

The utility of this 5 species community is demonstrated in the current and previous work by us, and its potential to test future ideas. For example, we have demonstrated how species become (mal)adapted after coevolutionary time [47] and how invaders and disturbances interact synergistically in their effect on resident diversity [58]. While the 5 species community as described only contains bacteria, we have also discovered three phages that each can infect one of the five species [59] and combined the model community with a broad host range plasmid which can be carried by all 5 species [60]. Moreover, we have used this community to demonstrate that fitness effects of plasmids shape the structure of bacteria– plasmid interaction networks [60]. Given its long-term stability and the distinct morphotypes of each species, this community is ideal to understand how microbial communities evolve and how this alters stability in the face of environmental stressors. Furthermore, its stability means this community is an ideal one to use to measure invasibility, or as a way to validate and improve Genome Scale Models (GSMs) that use genomic information to predict metabolic functioning and species interactions [61]. Environmental conditions such as primary substrate, pH, or temperature could be changed to see how well GSMs do under different scenarios.

## Data availability

All data and code for recreating the analyses and plots from the processed datasets is available on GitHub: https://github.com/padpadpadpad/five_species_community_paper. The assembled genomes have been submitted to NCBI and are available under the following accessions: *Achromobacter veterisilvae* AB1 (CP148753.1), *Ochrobactrum teleogrylli* AB1 (JBBHKQ000000000.1), *Pseudomonas fluorescens* AB1 (CP148752), *Stenotrophomonas sp.* AB1(2024)(JBBHKO000000000.1), and *Variovorax sp.* AB1(2024)(JBBHKP000000000.1). The raw sequencing of the original colony identification (Sanger sequencing) is available on the GitHub repository, and long-term colony identification (16s amplicon sequencing) will be made available on ENA. The raw short- and long-read whole genome sequencing data can be made immediately available on request, as can the code that creates the processed datasets.

## Acknowledgements

This work was funded by NERC. D.P. was also supported by a NERC Independent Research Fellowship (NE/W008890/1). E.H. was supported by a UKRI Future Leaders Fellowship (MR/V022482/1). D.S. was supported in part by grant MR/N0137941/1 for the GW4 BIOMED MRC DTP, awarded from the Medical Research Council.

