## Supplementary Information for "Characterising a stable five-species microbial community for use in experimental evolution and ecology"

**Table S1. The 46 bacterial strains used to establish the stable model community.**

| Code | Family | Genus |
|---|---|---|
| NMC6xB_4 | Alcaligenaceae | Achromobacter |
| NMC5_19 | Alcaligenaceae | Achromobacter |
| NMC1_18 | Comamonadaceae | Acidovorax |
| NMC6_14 | Microbacteriaceae | Agromyces |
| NMC2x_9 | Micrococcaceae | Arthrobacter |
| NMC1_7 | Micrococcaceae | Arthrobacter |
| NMC5_4 | Bacillaceae | Bacillus |
| NMC5_14 | Bacillaceae | Bacillus |
| NMC1B_24 | Alcaligenaceae | Bordetella |
| NMC5xB_11 | Alcaligenaceae | Bordetella |
| NMC5_15 | Caulobacteraceae | Brevundimonas |
| NMC6x_4 | Caulobacteraceae | Brevundimonas |
| NMC1B_19 | Alcaligenaceae | Candidimonas |
| NMC6_15 | Alcaligenaceae | Candidimonas |
| NMC2xB_19 | Burkholderiaceae | Cupriavidus |
| NMC1_10 | Burkholderiaceae | Cupriavidus |
| NMC1B_16 | Hyphomicrobiaceae | Devosia |
| NMC5_2 | Hyphomicrobiaceae | Devosia |
| NMC5x_2 | Flavobacteriaceae | Flavobacterium |
| NMC5x_7 | Flavobacteriaceae | Flavobacterium |
| NMC4_13 | Planococcaceae | Lysinibacillus |
| NMC3_4 | Planococcaceae | Lysinibacillus |
| NMC5B_21 | Microbacteriaceae | Microbacterium |
| NMC5_10 | Microbacteriaceae | Microbacterium |
| NMC4_4 | Brucellaceae | Ochrobactrum |
| NMC6B_13 | Cellulomonadaceae | Oerskovia |
| NMC2B_2 | Cellulomonadaceae | Oerskovia |
| NMC4_22 | Paenibacillaceae | Paenibacillus |
| NMC4x_9 | Paenibacillaceae | Paenibacillus |
| NMC6xB_20 | Rhodobacteraceae | Paracoccus |
| NMC5_5 | Rhodobacteraceae | Paracoccus |
| NMC5_17 | Sphingobacteriaceae | Pedobacter |
| NMC5_21 | Alcaligenaceae | Pigmentiphaga |
| NMC5_3 | Pseudomonadaceae | Pseudomonas |
| NMC6B_1 | Pseudomonadaceae | Pseudomonas |
| NMC6B_4 | Alcaligenaceae | Pusillimonas |
| NMC5_9 | Alcaligenaceae | Pusillimonas |
| NMC4B_5 | Rhizobiaceae | Rhizobium |
| NMC4B_1 | Nocardiaceae | Rhodococcus |
| NMC4B_16 | Nocardiaceae | Rhodococcus |
| NMC4B_3 | Rhodocyclaceae | Shinella |
| NMC3xB_22 | Staphylococcaceae | Staphylococcus |
| NMC5_6 | Xanthomonodaceae | Stenotrophomonas |
| NMC5_23 | Xanthomonodaceae | Stenotrophomonas |
| NMC1_21 | Comamonadaceae | Variovorax |
| NMC6_9 | Comamonadaceae | Variovorax |

**Table S2. Equations from calibration curves to estimate each species' CFU $uL^{-1}$ from optical density (OD) readings.**

| species | equation |
|---|---|
| *Achromobacter* sp. | (11676.OD - 291).16667 |
| *Ochrobactrum* sp. | (5548.OD - 67).16667 |
| *Pseudomonas* sp. | (546.OD - 20).16667 |
| *Stenotrophomonas* sp. | (56625.OD - 2246).16667 |
| *Variovorax* sp. | (955.OD - 30).16667 |
